# Convergent activity but divergent anxiety dimensions in male and female C57BL/6 mice across the open field and elevated plus maze

**DOI:** 10.64898/2026.09.13.751230

**Authors:** Guido Huisman, Lara S Caglayan, Marcelo Febo, Chengguo Xing, Adriaan W. Bruijnzeel

**Affiliations:** Department of Psychiatry, College of Medicine, University of Florida, Gainesville, FL, USA; Department of Medicinal Chemistry and Center for Natural Products, Drug Discovery and Development (CNPD3), College of Pharmacy, University of Florida, Gainesville, FL, USA

**Keywords:** C57BL/6, mice, sex differences, open field, elevated plus-maze, anxiety-like behavior, locomotor activity, habituation, hierarchical clustering

## Abstract

Anxiety disorders are among the most common psychiatric illnesses, with a significantly higher prevalence in females than in males. The open field (OF) and the elevated plus maze (EPM) are the two assays used most often to model anxiety-like behavior in mice. It is usually assumed that they index a common underlying trait, yet few studies have investigated this in a large group of male and female mice using an advanced analytical approach. The goal of these studies was to determine sex differences in locomotor activity and anxiety-like behavior in the OF and the EPM, and to establish which behavioral parameters correlate between the tests. Adult male and female C57BL/6 mice (n = 61; 29 male, 32 female) were given a single 15 min session in the OF, analyzed as a whole and in three consecutive 5 min blocks, and a single 5 min EPM trial. Males and females did not differ on any OF measure over the cumulative 15-minute session. Locomotion and stereotypy declined over time with females habituating faster than males while center exploration did increase. In the EPM, females made more closed-arm entries than males while open-arm exploration was identical. Every significant correlation between the two tests involved an activity measure. All four pairs formed between OF and EPM activity parameters were significant whereas none of the six pairs formed between OF center parameters and EPM open-arm parameters were significant. Hierarchical clustering revealed the same structure, placing the activity measures of both tests in one cluster, the EPM open-arm measures in a second, and the two OF center measures in a third. The cross-test activity correlation was strongest during the initial five minutes of the OF and weakened as the session progressed. These studies indicate that the OF and EPM yield convergent measures of activity, but divergent measures of anxiety-like behavior.

## 1 Introduction

Anxiety disorders are one of the most prevalent classes of psychiatric illnesses globally and impose a significant psychosocial, economic, and healthcare burden (Bandelow and Michaelis, 2015). Epidemiological studies have consistently demonstrated profound sex disparities in anxiety disorders. Women exhibit a higher lifetime prevalence of generalized anxiety disorder, panic disorder, specific phobias, agoraphobia, and post-traumatic stress disorder compared to men, and the symptoms are often more severe and of longer duration in women (McLean et al., 2011; Altemus et al., 2014; Kessler et al., 1994). Despite this overrepresentation of females in clinical populations, historically preclinical studies have relied upon male rodents (Beery and Zucker, 2011). This focus on males in animal studies has limited the translational validity of preclinical models and potentially obscured sexually dimorphic neurobiological mechanisms that govern threat appraisal, stress responsivity, and emotional regulation (Cahill, 2006).

During the last decade, a paradigm shift has underscored the necessity of integrating sex as a biological variable in biomedical research (Clayton and Collins, 2014). Neuroimaging and transcriptomic studies have revealed significant structural, functional, and molecular differences between the male and female brain (Liu et al., 2020a; Wingo et al., 2023). These sex differences are highly pronounced within stress-responsive corticolimbic circuits, including the medial prefrontal cortex, hippocampus, amygdala, and the bed nucleus of the stria terminalis (BNST) (Liu et al., 2020b; Turano et al., 2019). Therefore, there is an urgent need to characterize sex differences in animal models of psychiatric disorders.

To investigate the neurobiology of emotionality and screen for potential anxiolytic compounds, most studies rely on unconditioned behavioral assays. These tasks are based on the innate approach-avoidance conflict between a rodent’s drive to explore novel environments and its aversion to exposed spaces (Milner and Crabbe, 2008). Due to genetic advances, mice have largely replaced rats as the premier model organism for neuroscience studies (Milner and Crabbe, 2008; Ellenbroek and Youn, 2016). Two of the most widely used and extensively validated paradigms for evaluating anxiety-like behavior in rodents are the open field (OF) test and the elevated plus-maze (EPM) (Hogg, 1996; Prut and Belzung, 2003; Tan et al., 2019).

The OF test was originally developed to evaluate emotionality through ambulation and assess approach-avoidance conflict by measuring the animal’s willingness to move away from the protective peripheral walls (thigmotaxis) and explore the center of the arena (Hall, 1934; Prut and Belzung, 2003). The OF test is somewhat stressful, as indicated by increased corticosterone levels, and in this test, reduced center exploration is traditionally interpreted as a measure of heightened anxiety-like behavior (Kas et al., 2005; Prut and Belzung, 2003). However, despite this interpretation, classical anxiolytics such as benzodiazepines fail to increase center exploration in the open field test in mice, including in C57BL/6J mice (Thompson et al., 2015; Birkett et al., 2011). The EPM relies on the rodent’s innate fear of heights and open spaces (agoraphobia). The apparatus consists of two enclosed, safe arms and two exposed, open arms that are elevated above the ground (Pellow et al., 1985). The EPM has been extensively validated with clinically effective anxiolytics; for example, benzodiazepines such as diazepam and chlordiazepoxide selectively increase the percentage of entries and time spent in the open arms, whereas anxiogenic stimuli decrease these measures (Pellow et al., 1985; Braun et al., 2011). Furthermore, other anxiolytic drugs such as opioids also decrease anxiety-like behavior in the EPM (Bruijnzeel et al., 2022).

The OF and EPM are commonly used as parallel measures of a unitary “anxiety” construct. Therefore, if both the OF and the EPM measure a common latent dimension of trait anxiety, an individual animal displaying a high level of anxiety in the OF (e.g., profound center avoidance) should theoretically exhibit high anxiety in the EPM (e.g., profound open-arm avoidance). However, some empirical data contradict this assumption. Previous preclinical studies have shown that behavioral correlations across different ethological paradigms can be variable (Ramos, 2008).

Conflicting findings have been reported regarding sex differences in anxiety tests. The expression of sex differences in the OF and EPM is sensitive to experimental conditions, including strain, age, lighting intensity, and pre-test handling (Knight et al., 2021). For instance, a comprehensive evaluation in drug-naïve adult Wistar rats demonstrated robust sex differences across virtually all parameters of the OF and EPM, with females consistently displaying less anxiety-like behavior than males (Knight et al., 2021). Conversely, numerous other studies utilizing rats, or various mouse strains such as the widely used C57BL/6J, have reported either no sex differences or heightened anxiety-like behavior in females (Milner and Crabbe, 2008; Knight et al., 2021; Tucker and McCabe, 2017; Johnston and File, 1991). These discrepancies are often exacerbated by insufficient statistical power and differences in baseline locomotor activity (Knight et al., 2021). Thus, to provide insight into sex differences, studies require rigorous analytical approaches that account for general locomotion and the temporal dynamics of behavior.

Given the multidimensionality of anxiety and the inconsistencies regarding sex differences in anxiety tests, the present study was designed to determine whether behavioral parameters in the OF and the EPM correlate with one another in drug-naïve adult mice, and whether the two tests reveal the same sex differences. Because a 15 min OF session is not behaviorally uniform, the session was analyzed both as a whole and in three consecutive 5 min blocks, which allows the cross-test relationship to be followed as the animal habituates to the OF arena.

## 2 Methods

### 2.1 Animals

Adult male and female C57BL/6NCrl mice (8 weeks of age; N = 32 per sex) were purchased from Charles River (Raleigh, NC). Because of fighting-related injuries, three males were not included in the behavioral tests, resulting in a final sample of 61 mice (29 males, 32 females). During the one-week acclimatization period the mice were handled and weighed on several days, and body weight on the day of open field testing was 23.9 ± 0.3 g in the males and 18.9 ± 0.2 g in the females.

Mice of the same sex were socially housed in groups of four, unless a cage mate was lost, in a climate-controlled vivarium (22°C with 50% humidity) with a 12 h light-dark cycle (lights on between 07:00–19:00). Food and water were available ad libitum. The experimental protocols were approved by the University of Florida Institutional Animal Care and Use Committee (IACUC). All experiments were performed in accordance with relevant IACUC guidelines and regulations and in compliance with ARRIVE guidelines 2.0 (Animal Research: Reporting of In Vivo Experiments).

### 2.2 Experimental design and behavioral testing

All behavioral testing was conducted during the light phase between 13:00 and 17:00. To prevent the influence of olfactory cues from previous subjects, all testing arenas were cleaned with Nolvasan (chlorhexidine diacetate) between each animal. Following the one-week vivarium habituation period, each mouse was given a single 15 min session in the OF. Two days later, the animals were tested on the EPM for 5 min. No injections, surgical interventions or experimental diets were administered before or during the behavioral testing.

### 2.3 Open field test

Locomotor activity and time in center were measured using an automated rodent activity monitoring system (AccuScan Instruments, Columbus, OH) under low light conditions (Febo et al., 2024; Chellian et al., 2024). The activity cages were made from clear Plexiglas (40 × 40 × 30 cm; L × W × H) with an outer steel frame and were equipped with infrared emitting and receiving panels. Each panel contained 16 evenly spaced infrared beams (2.5 cm apart) crossing the length and width of the cage at a height of 2 cm, which enabled detection of horizontal movements. The arena was divided into two distinct zones: an outer (margin) zone 7.5 cm wide and a center zone of 25 × 25 cm. Patterns of sequential and repetitive infrared beam interruptions were recorded and transmitted to a programmable counter, which sorted the beam counts and sent the data to a PC running VersaMax™ software. The activity data were categorized into total distance traveled, center distance, time in the center, and stereotypy count. The percentage of center time (center time / session duration × 100) and the percentage of center distance (center distance / total distance × 100) were calculated. Activity data were analyzed for the full session and in 5 min bins. Between each trial, the OF was cleaned with a Nolvasan solution.

### 2.4 Elevated plus-maze test

The mouse EPM apparatus consisted of two opposing open arms (31 × 6 cm; L × W) and two opposing closed arms (31 × 6 × 15 cm, L × W × H) extending from a center platform (6 × 6 cm; L × W), elevated 41 cm above the floor as described before (Wang et al., 2016). Testing was conducted in a quiet, dimly lit room (75 lx). At the beginning of each 5-min test, mice were placed in the center of the apparatus facing an open arm and allowed to freely explore. The apparatus was cleaned with a Nolvasan solution between animals. Animal behavior was recorded with a ceiling-mounted camera and analyzed automatically via center-point detection using EthoVision XT 19 software (Noldus Information Technology, Leesburg, VA) as described before (Tan et al., 2019; Qi et al., 2016). The EPM was divided into five zones: two open arms, two closed arms, and a center zone. The software automatically calculated the total distance traveled, as well as the duration and number of entries into each respective zone. From these raw parameters, the percentage of open-arm entries (open-arm entries / total arm entries × 100) and the percentage of time spent on the open arms (time on open arms / total time on arms × 100) were calculated.

### 2.5 Statistical analysis

Behavioral parameters in the OF and the EPM were analyzed with a one-way ANOVA with Sex as a between-subjects factor. Behavior across the three 5 min blocks of the OF session was analyzed with a two-way mixed ANOVA with Block (minutes 0–5, 5–10 and 10–15) as the within-subjects factor and Sex as the between-subjects factor. Bonferroni post-hoc comparisons were conducted when there were significant Block × Sex interactions.

Pearson’s correlation coefficients (r) were calculated within and between tests, first with the sexes combined (n = 61) and then separately in males (n = 29) and females (n = 32). P values (two-sided t test) were calculated to determine whether the correlation coefficients were significant and were corrected for multiple comparisons with the Holm–Bonferroni procedure. The Holm family comprised the 20 cross-test pairs (four OF measures × five EPM measures) within a given analysis window in a given group. Within-test correlations were treated as descriptive and were not subjected to multiple comparison correction. A sensitivity analysis indicated that with n = 61 and α = .05 (two-tailed, uncorrected), the study had 80% power to detect correlations of r ≥ 0.35. Spearman’s correlations were also calculated to confirm the results were independent of the distribution of the data. Whether the correlation between the OF and the EPM differed between the sexes was tested with Fisher’s r-to-z transformation.

Partial correlations that controlled for OF total distance were calculated to determine whether cross-test relationships were independent of activity. For the hierarchical cluster analysis, correlation coefficients were converted to distances as d = √(2(1 − r)), and the resulting distance matrix was clustered with Ward’s method (Ward Jr, 1963; Knight et al., 2021). Clusters were defined by a conventional cut at 70% of the maximum linkage height.

To avoid inflating the correction family with redundant measures, both parameter sets were reduced before the analysis using an a priori redundancy criterion. When two parameters correlated at |r| ≥ 0.85, or one was an arithmetic transform of another, a single representative was kept. In the OF, margin time and center time were omitted (margin time + center time = session duration and therefore both are exact linear functions of percentage center time) and horizontal activity was omitted as redundant with total distance (r = +0.93), leaving four parameters for the correlation analysis: percentage center time, percentage center distance, total distance and stereotypy count. In the EPM, open-arm duration was omitted (r = +0.99 with the percentage of time on the open arms), closed-arm duration was omitted (r = −0.955 with the percentage of time on the open arms) and center entries were omitted (r = +0.91 with distance traveled on the EPM), leaving five parameters for the correlation analysis: the percentage of open-arm entries, the percentage of time on the open arms, open-arm entries, closed-arm entries and distance traveled. The reduced set of five was used for the correlation, partial correlation and clustering analyses, giving 20 cross-test correlations in each analysis window in each group. For descriptive purposes the parameters were grouped a priori as activity measures (OF total distance and stereotypy count; EPM distance traveled and closed-arm entries) and anxiety-related measures (OF percentage center time and percentage center distance; EPM percentage of open-arm entries, percentage of time on the open arms, and open-arm entries). These groupings define four activity pairs and six anxiety pairs among the 20 cross-test correlations.

The OF and EPM heatmaps were generated in Python. For the OF, the raw beam-coordinate files exported by the VersaMax system were used. For each animal the interval between successive samples was assigned to the photobeam cells the animal occupied at that sample and expressed as a percentage of the session. For the EPM, the raw x,y coordinates exported by EthoVision were binned on a spatial grid and expressed as a percentage of the tracked samples and smoothed with a Gaussian kernel (σ = 1 bin). The maps were then averaged across animals within each sex and displayed on a color scale.

Analyses of variance were performed in IBM SPSS Statistics Version 32. Correlation, partial correlation and clustering analyses were performed in Python (Google Colab) using pandas (McKinney, 2010), SciPy (Virtanen et al., 2020), statsmodels (Seabold and Perktold, 2010) and pingouin (Vallat, 2018). Figures were prepared in matplotlib and GraphPad Prism Version 11. Statistical significance was set at p < 0.05.

## 3 Results

### 3.1 Sex differences in the open field

No sex difference was detected in any OF measure over the 15 min session. Males and females spent a comparable proportion of the session in the center zone (F(1,59) = 0.50, p = 0.48) and covered a comparable proportion of their distance there (F(1,59) = 0.41, p = 0.53). Total distance traveled (F(1,59) = 0.07, p = 0.79) and stereotypy count (F(1,59) = 1.70, p = 0.20) also did not differ (Figure 1 and 2, Table S1).

**Figure 1.**
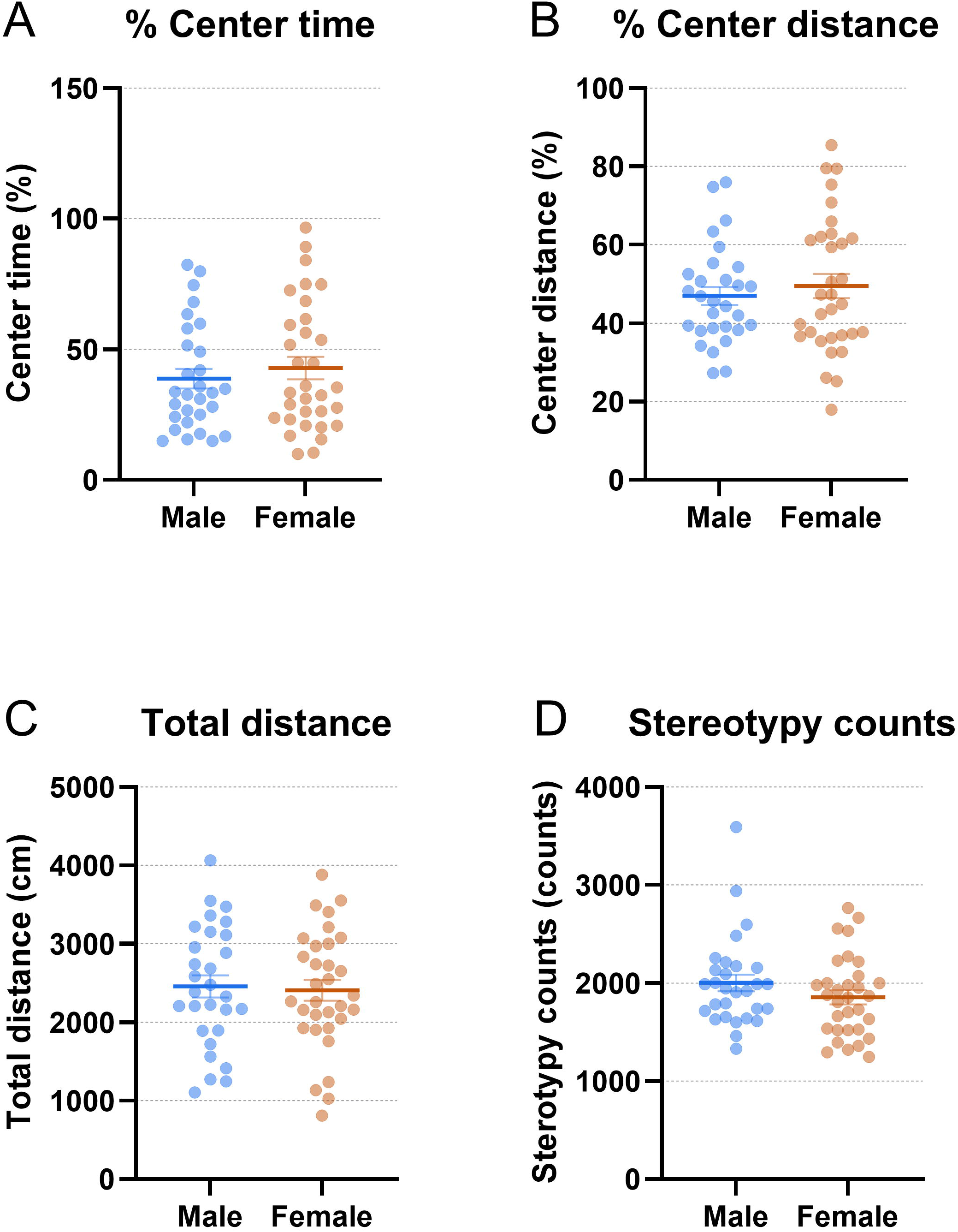
Male and female mice display similar behaviors in an open field session. Percentage of the session spent in the center zone (A), percentage of distance traveled in the center zone (B), total distance traveled (C) and stereotypy counts (D), over the 15 min OF session. Points are individual animals with the group mean ± SEM; males n = 29, females n = 32.

**Figure 2.**
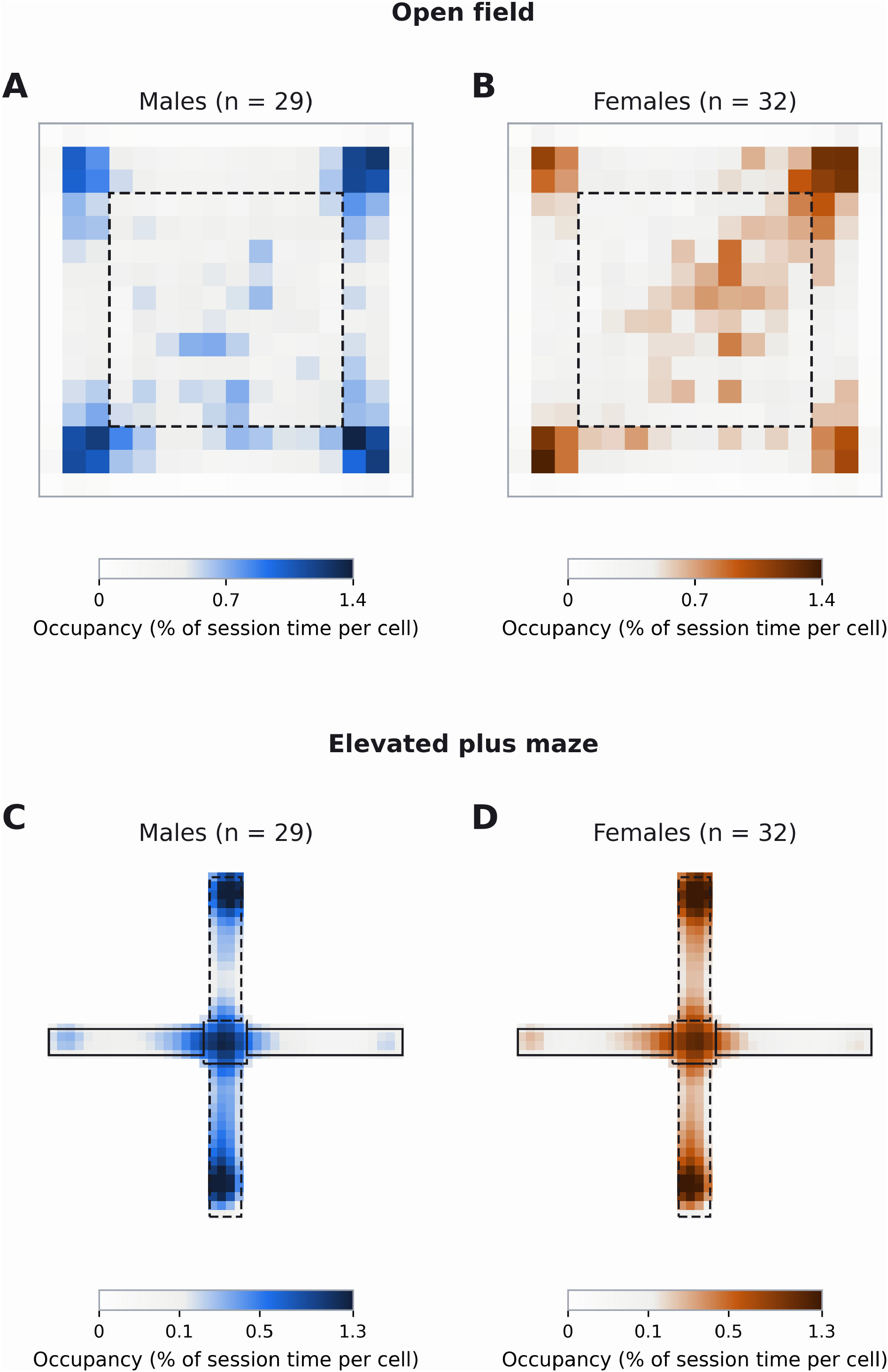
Open field and elevated plus maze heatmaps. Spatial distribution of time spent by the mice in the different parts of the OF and the EPM. Mean occupancy across the 15-min baseline OF session for males (A, n = 29) and females (B, n = 32), and across the 5-min EPM for males (C, n = 29) and females (D, n = 32). Color indicates the percentage of session time spent in each cell of the apparatus. The dashed square in A and B marks the open field center zone. In C and D the solid outlines mark the open arms and the dashed outlines the closed arms and the center platform. Open field panels use a linear color scale spanning 0 to 1.4% of session time per cell; elevated plus maze panels are smoothed and use a square-root color scale spanning 0 to 1.3%.

### 3.2 Within-session habituation of activity, and its absence for center exploration

Locomotor activity (Block F(2,118) = 55.49, p < 0.001) and stereotypies (Block F(2,118) = 17.89, p < 0.001) declined over the 15 min OF test (Figure 3). Percentage center time increased over time (Block F(2,118) = 4.37, p = 0.015), and percentage center distance also increased (Block F(2,118) = 16.26, p < 0.001). Percentage center time or distance did not show a Block × Sex interaction (p = 0.66 and p = 0.65).

**Figure 3.**
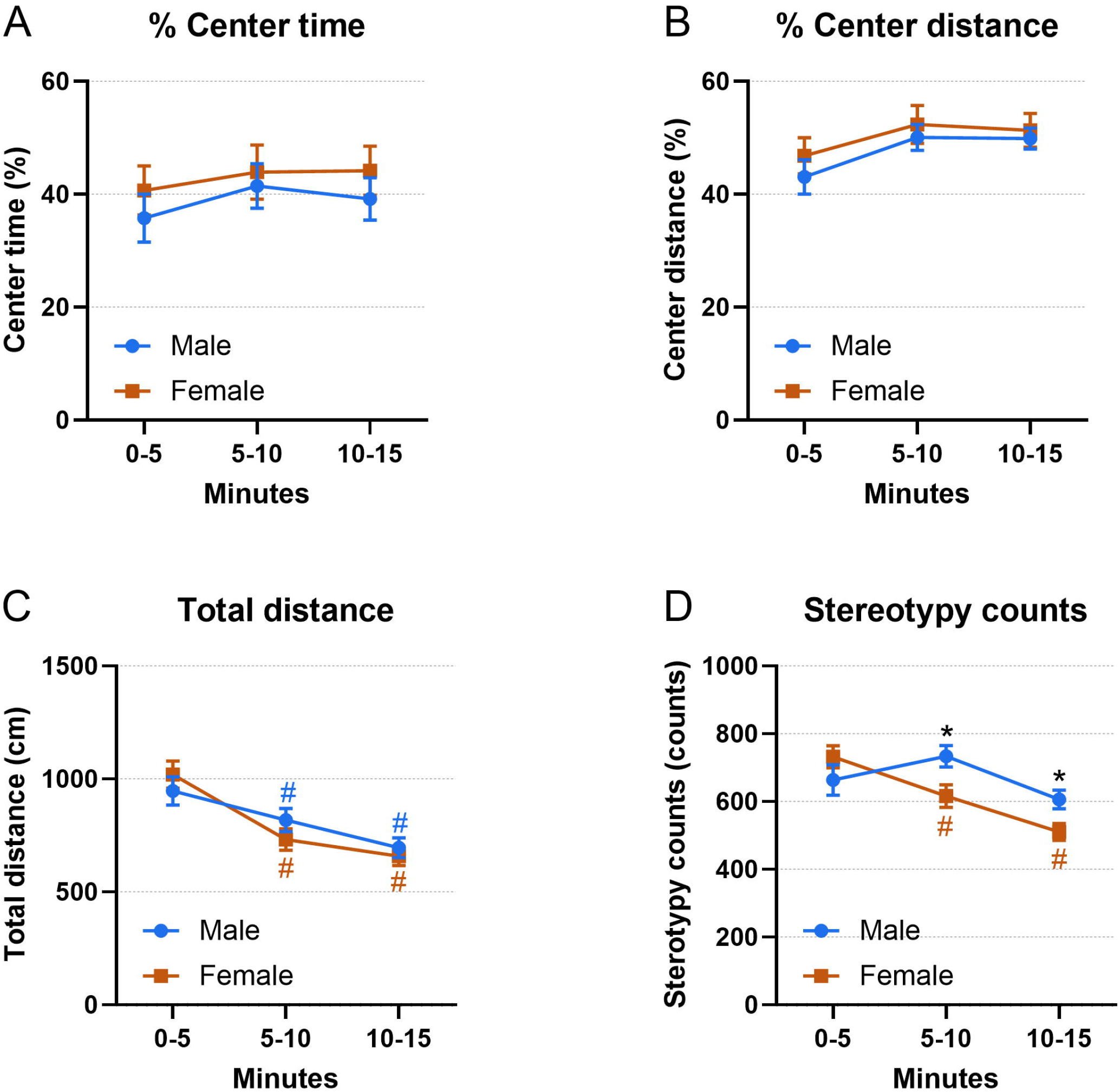
Sex-dependent habituation of locomotor activity in the open field. Percentage of time spent in the center zone (A), percentage of distance traveled in the center zone (B), total distance traveled (C) and stereotypy counts (D), over minutes 0–5, 5–10 and 10–15. Symbols are group means ± SEM for males (n = 29) and females (n = 32). Locomotion and stereotypy declined across the session while center exploration increased. Pound sign (#) indicates that a block differs from that same sex’s own first block and asterisk (*) marks a difference between the sexes at that block. Both symbols denote corrected p < 0.05.

Females habituated faster than males as indicated by the fact that the Block × Sex interaction was significant for total distance (F(2,118) = 3.75, p = 0.026) and for stereotypy count (F(2,118) = 8.17, p < 0.001), and neither measure showed a main effect of Sex (total distance, F(1,59) = 0.07, p = 0.79 and stereotypy count, F(1,59) = 1.70, p = 0.20).

Post-hoc comparisons showed that both sexes reduced the distance traveled from the first block onwards (all corrected p ≤ 0.023). Females reduced their stereotypy count (corrected p ≤ 0.009 between every pair of blocks) but males did not differ from their own first block at either later block (corrected p = 0.36 and p = 0.38).

Post-hoc tests were also conducted to examine sex differences within blocks. Males and females did not differ in the first five minutes (stereotypy corrected p = 0.67, total distance corrected p = 1.00), and males displayed more stereotypies by minutes 5–10 (F(1,59) = 6.25, corrected p = 0.046) and minutes 10–15 (F(1,59) = 6.53, corrected p = 0.040) compared to females. For total distance, the post hoc tests did not reveal significant differences between the males and the females (all corrected p ≥ 0.67).

### 3.3 Sex differences in the elevated plus maze

Female mice made more closed-arm entries than males (F(1,59) = 10.54, p = 0.0019)(Figure 2 and 4, Table S2). Open-arm exploration did not differ between the sexes on any measure: open-arm entries (F(1,59) = 0.11, p = 0.74), open-arm duration (F(1,59) = 0.32, p = 0.57), the percentage of open-arm entries (F(1,59) = 3.50, p = 0.067) and the percentage of time spent on the open arms (F(1,59) = 0.73, p = 0.40). There was a non-significant trend towards the females traveling a greater distance on the EPM (F(1,59) = 3.24, p = 0.077), and there were also no sex differences in center entries (F(1,59) = 0.46, p = 0.50) or closed-arm duration (F(1,59) = 0.77, p = 0.38). Females were therefore somewhat more active on the maze than males, and this additional activity was directed into the enclosed arms.

**Figure 4.**
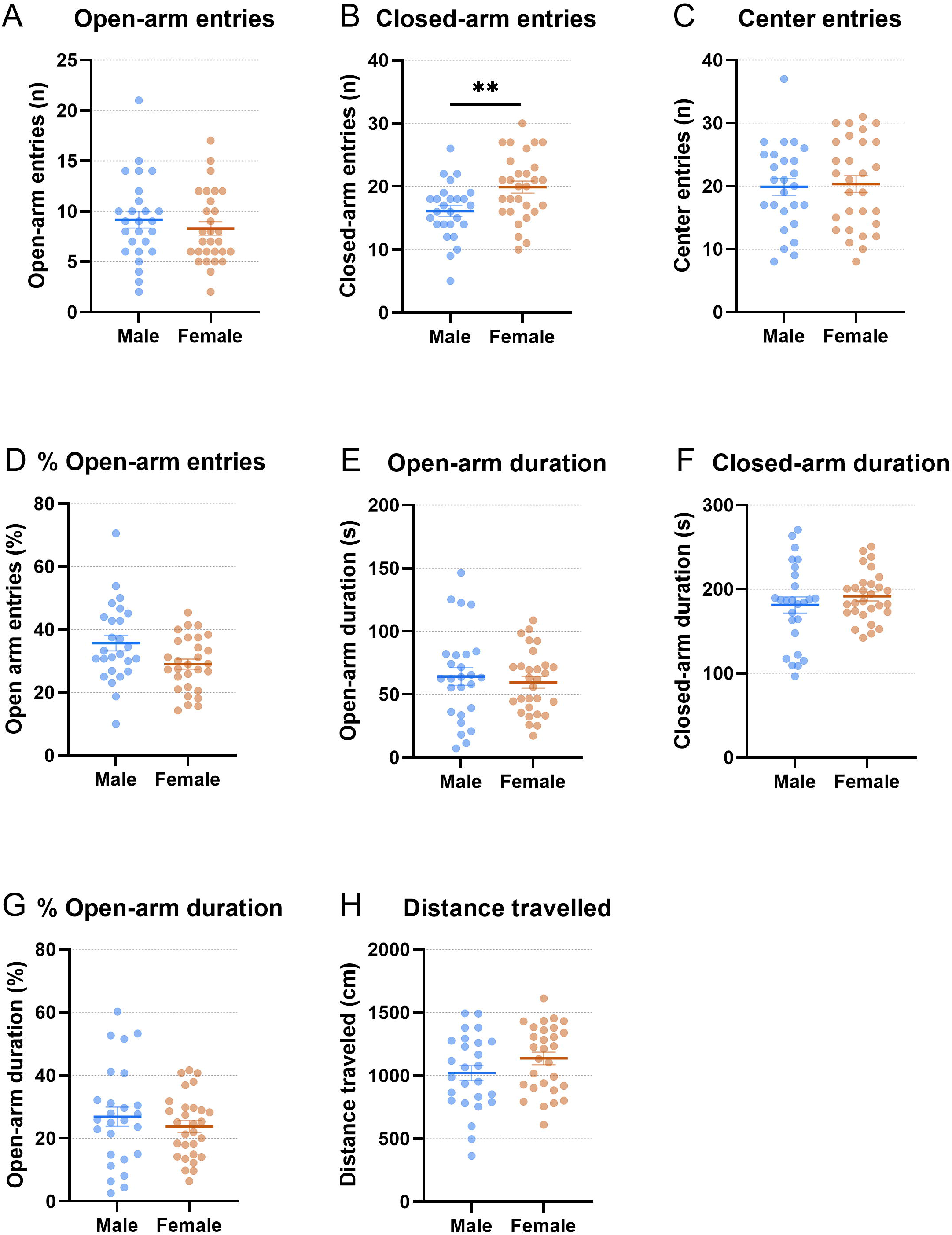
Female mice demonstrate increased closed-arm entries without altered open-arm exploration in the elevated plus-maze. Eight behavioral parameters recorded in the elevated plus maze (A–H). Points are individual animals with the group mean ± SEM; males n = 29, females n = 32. Asterisks denote a sex difference: ** p < 0.01.

### 3.4 Open field and elevated plus maze cross-test correlations

Within-test correlations were strong in both tests, whereas correlations between the tests were modest and largely confined to a single behavioral dimension (Figure 5). Of the 20 cross-test pairs computed over the whole 15 min OF session, five were significant, and every one of them involved an activity measure in the OF (total distance or stereotypy count). All four pairs formed between the two OF activity measures and the two EPM activity measures (distance traveled, closed-arm entries) were significant, with a mean |r| of 0.332. The fifth was OF stereotypy count with EPM open-arm entries (r = +0.262, p = 0.041). Two survived Holm correction across the family of 20: OF stereotypy count with EPM distance traveled (r = +0.389, uncorrected p = 0.0020) and with EPM closed-arm entries (r = +0.389, uncorrected p = 0.0020), both with an adjusted p of 0.039. Both were reproduced by Spearman’s rank correlation (ρ = +0.380, p = 0.0025 and ρ = +0.367, p = 0.0037), confirming that they do not depend on distributional assumptions.

**Figure 5.**
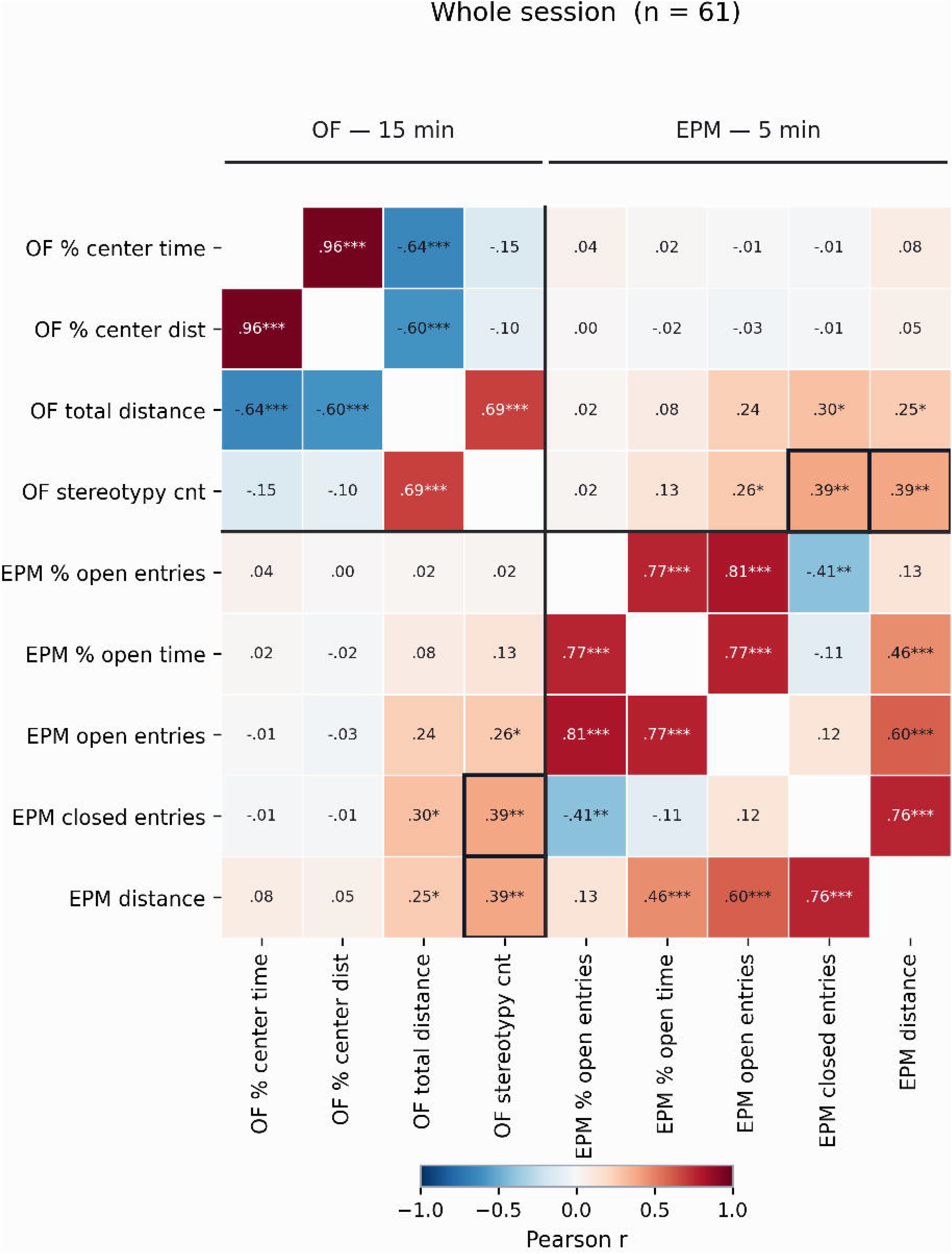
Cross-test correlations reveal convergent activity metrics but no relationship between anxiety-like parameters. Correlation matrix for the pooled sample (n = 61) computed from the whole 15 min OF session against the 5 min plus-maze trial. Every significant cross-test correlation involves an activity measure in the OF. There were no significant cross-test correlations between OF center measures and plus-maze open-arm measures. Cells show Pearson correlation coefficients; the color scale runs from r = −1 (blue) through r = 0 (white) to r = +1 (red). Asterisks denote uncorrected two-tailed p values: * p < 0.05, ** p < 0.01, *** p < 0.001. Bold outlines mark cells that survive Holm correction.

Not one of the six pairs formed between an OF center measure (percentage center time, percentage center distance) and an EPM open-arm measure (the percentage of open-arm entries, the percentage of time on the open arms, open-arm entries) approached significance. The mean |r| across those six pairs was 0.019 and the largest was 0.043; the correlation between OF percentage center time and the EPM percentage of open-arm entries was r = +0.043 (p = 0.74), and Spearman’s rank correlation provided the same outcome (ρ = +0.105, p = 0.42). Controlling for OF total distance did not reveal a new relationship that had been masked by general activity (partial r = +0.073, p = 0.58 for percentage center time; +0.016, p = 0.90 for percentage center distance). The same pattern was present in each sex analyzed separately, with three significant activity pairs in males and two in females and no significant anxiety pair in either, and Fisher’s r-to-z also indicated that the cross-test correlation did not differ between the sexes (all p ≥ 0.12) (Figure S1).

### 3.5 The cross-test activity correlation weakens across the session

Because activity habituated within the session, the cross-test correlations were recalculated separately for each 5 min block (Figure 5, S2). The activity coupling was strongest at the start of the session and weakened thereafter: three of the four activity pairs were significant in minutes 0–5 (mean |r| = 0.359; the strongest, OF stereotypy count with EPM closed-arm entries, was r = +0.466, uncorrected p = 0.0002, Holm-adjusted p = 0.003), one in minutes 5–10 (mean |r| = 0.234) and none in minutes 10–15 (mean |r| = 0.199) (Figure 6A). The anxiety pairs remained at zero throughout, with mean |r| values of 0.029, 0.022 and 0.036 across the three blocks and no pair significant in any block (Figure 6B). The correlations between the two tests are therefore not only restricted to activity but are also transient. The correlations are strongest during the first 5 min immediately after the mice are placed in the OF.

**Figure 6.**
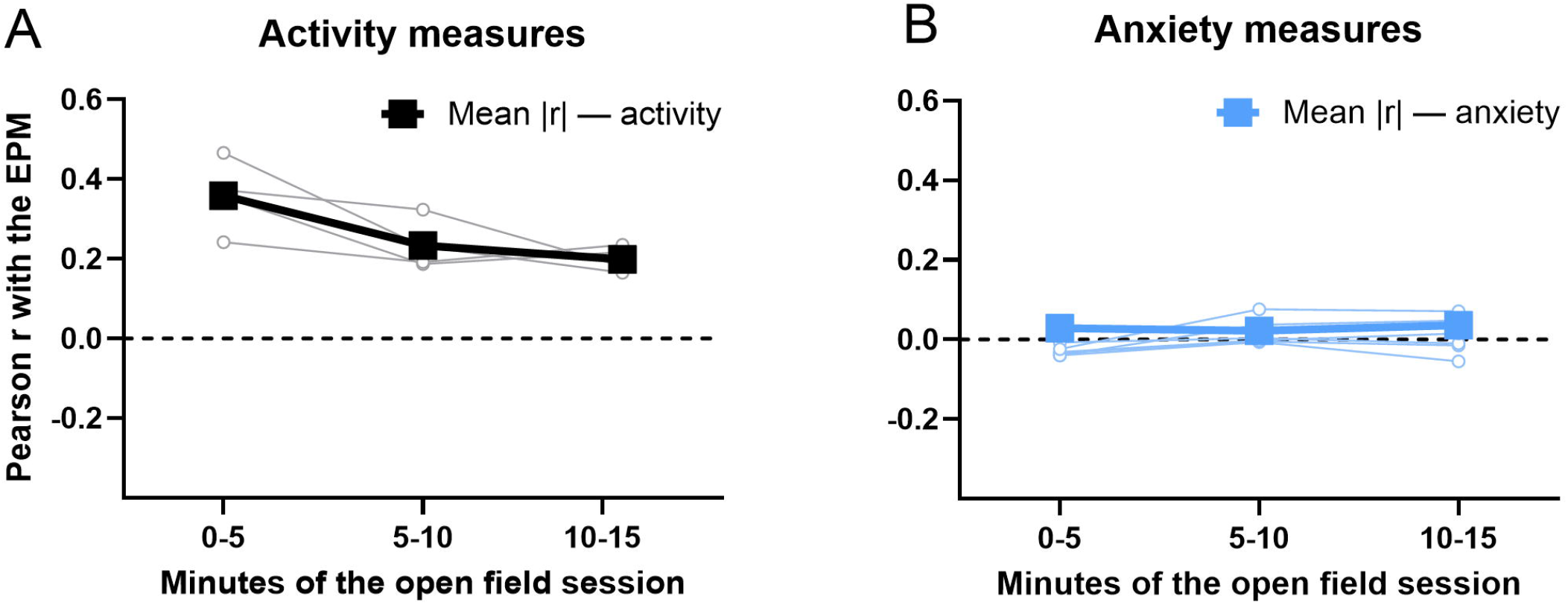
The cross-test activity correlation is strongest during initial novelty exposure and declines over the open-field session. The cross-test activity correlation is strongest during initial novelty exposure and declines over the OF session. Pearson correlations with the EPM plotted against the block of the OF session in which the OF measure was scored. The thin lines are individual pairs and the thick line is the mean absolute correlation within each panel. Correlations are shown for the four pairs formed between OF activity measures (total distance, stereotypy count) and the two plus-maze activity measures, distance traveled and closed-arm entries (A), and for the six pairs formed between OF center measures (center time, center distance) and the three plus-maze open-arm measures, the percentage of open-arm entries, the percentage of time on the open arms, and open-arm entries (B).

### 3.6 Hierarchical clustering

Ward’s clustering of the correlation distance matrix, cut at 70% of the maximum linkage height, separated the nine behaviors into three clusters (Figure 7). The first contained OF total distance and stereotypy count together with EPM distance traveled and closed-arm entries; the second contained the three EPM open-arm measures; and the third contained the two OF center measures alone. The only cluster combining behaviors from both tests was therefore the activity cluster, and neither OF center measure grouped with any EPM open-arm measure. The clustering confirms that the OF and EPM share a locomotor dimension (Figure 6).

**Figure 7.**
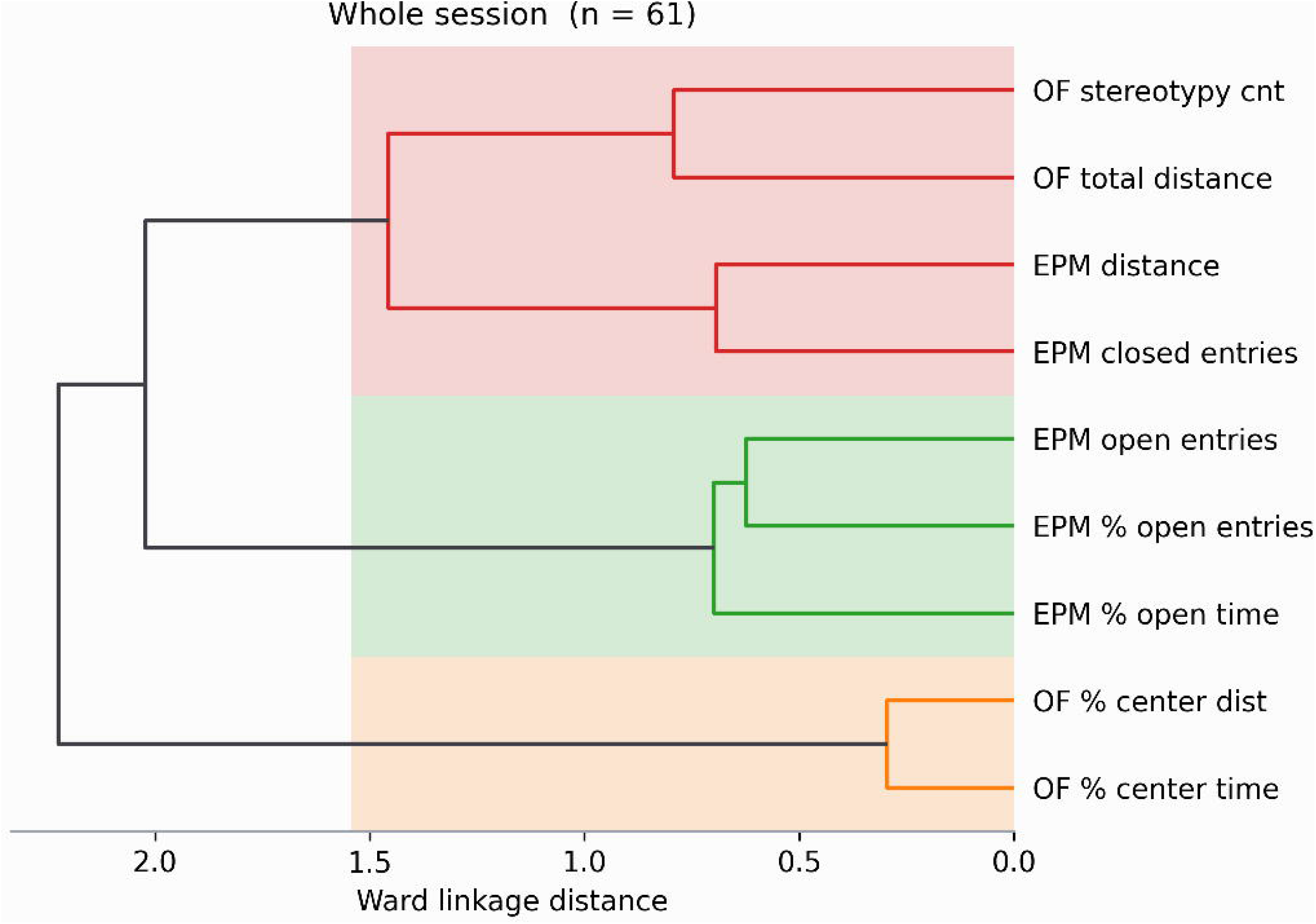
Hierarchical clustering separates a shared locomotor dimension from independent anxiety domains. Hierarchical clustering was used to group the four OF and five EPM parameters based on how strongly they correlate. Shading indicates the clusters obtained by cutting the tree at 70% of the maximum linkage height. The general activity measures were the only variables from both tests to group together into a single cluster. The OF center measures formed their own isolated cluster which was completely separate from all EPM open-arm measures.

## 4 Discussion

The present study compared the behavior of the same adult drug naïve C57BL/6N male and female mice in the OF and EPM. Four findings emerged. Sex differences were absent over the whole session from every OF measure and were confined in the EPM to activity directed into the closed arms. Locomotion habituated within the OF session while center exploration did increase, and the rate of that habituation differed between the sexes. Locomotor activity and stereotypies habituated faster in females than males with the males showing more stereotypies toward the end of the session. Finally, every significant correlation between the two tests involved an activity measure, with no relationship between OF center exploration and EPM open-arm exploration. Hierarchical clustering independently recovered this exact same structural division.

In the EPM, male and female mice had similar open-arm entries and open-arm durations. The female mice had a significantly higher number of closed-arm entries than males, however, although the females traveled a greater distance on the EPM this did not reach statistical significance. The lack of sexual dimorphism in anxiety parameters in our study is consistent with a previous study using C57BL/6J mice, which also found that male and female mice spend a similar amount of time in the open arms during their initial exposure to the EPM (Tucker and McCabe, 2017). Furthermore, in line with our finding of no significant difference in total distance, Tucker and McCabe reported that males and females travel similar distances during their first EPM exposure (Tucker and McCabe, 2017). In the present study, the females had more closed arm entries than the males. Increased closed-arm entries are traditionally viewed as a measure of generalized locomotor activity, as they are less affected by the approach-avoidance conflict than open-arm entries (Pellow et al., 1985). Therefore, a somewhat reductive interpretation would be that females are just more active on the maze. However, this increased activity is directed into the enclosed, “safe” arms rather than reflecting an indiscriminate increase in exploration across the entire apparatus. Alternatively, this behavioral profile may represent an active, hyper-vigilant coping strategy characterized by continuous scanning and patrolling of the safe environment. The lack of a sex difference in the open arms, paired with the higher activity in the closed arms, led to the percentage of open-arm entries being somewhat lower in females. This may be consistent with a more anxious or hyper-reactive phenotype in the females.

To get a better impression of the change in behavior in the OF over time during the 15-minute session we also analyzed the behavior in 5-minute blocks. Locomotion fell by roughly a third and stereotypy by a fifth across fifteen minutes, providing evidence for substantial intrasession habituation. Center exploration did not follow this pattern. The percentage center time and percentage center distance rose across the three blocks. Therefore, the decline in activity was not due to an increase in avoidance of the center. Interestingly, the rate of habituation was sexually dimorphic. Females reduced their distance traveled and stereotypies more than males and this led to sex differences in stereotypies that were absent in the opening five minutes and present by minutes 5–10 and 10–15. This highlights the importance of temporal binning in studies that investigate sex differences in rodents.

It remains a subject of ongoing debate whether ethological assays such as the OF and EPM measure a unified latent psychological construct of anxiety, or distinct dimensional features of emotionality (Ramos, 2008). The present data provide detailed insight into the relationship between behavioral phenotypes across these paradigms. Specifically, the assays demonstrated robust concordance regarding general motor activity. All four correlational pairs generated between OF locomotor measures (total distance, stereotypy count) and EPM activity indices (distance traveled, closed-arm entries) were significant. This cross-test locomotor convergence is consistent with prior work showing that OF ambulation correlates with arm visits in mice, although those correlations were obtained across two strains (C57BL/6J and BALB/c) rather than within one (Lalonde and Strazielle, 2008). However, while their study found no cross-test relationship for stereotypies, our data demonstrate that both ambulation and stereotypies share an activity dimension with enclosed-arm exploration. Furthermore, we extend these findings by showing that this activity relationship is dynamic. The correlation peaks during initial novelty and weakens across the session, and the rate of habituation is sexually dimorphic.

In contrast to the activity measures, the anxiety measures showed no relationship across the two tests. None of the six pairs formed between OF center measures and plus-maze open-arm measures approached significance, and the largest was only r = +0.043. With 61 animals the study had 80% power to find a correlation of 0.35 or larger, and therefore weak associations between the OF and EPM parameters cannot be ruled out. We can, however, rule out strong associations like those observed for the activity measures. For example, OF stereotypy correlated with EPM distance and with closed-arm entries at r = +0.389, which is above the threshold the study was powered to detect. In contrast, nothing approaching such a strong correlation was detected for the anxiety measures.

The activity measures therefore act as a positive control, showing that the design detected cross-test agreement of that size in this experiment. No such agreement was found for center and open-arm exploration, which points to a real dissociation rather than to a limitation of the study. These assays therefore do not appear to capture a unified, stable trait for anxiety-like behavior, which may instead reflect responses that are more state- and context-dependent.

To better understand the relationships among the OF and EPM parameters, hierarchical clustering was applied. This method groups variables by shared variance and provides a data-driven map of behavioral architecture (Knight et al., 2021). The tree was cut at a conventional height, 70% of its maximum linkage distance. Three groups emerged at that level, and the number reflects the structure of the correlations rather than a value chosen beforehand. The activity measures were the only cluster containing behaviors from both tests, while the two OF center measures formed a cluster of their own, separate from every EPM open-arm measure. Because this analysis uses no significance threshold and no correction for multiple comparisons, its agreement with the correlation matrix supports the conclusion that, under the present testing conditions, the two assays capture behavioral domains that overlap on general activity but are otherwise largely independent (Ramos, 2008). This is consistent with principal component analyses in C57BL/6 mice showing that EPM parameters such as arm entries load onto a general activity component that is independent of the components representing avoidance of specific test environments (Clément et al., 2007).

These data provide useful context for the design of preclinical behavioral studies. The practice of relying on a single testing environment to determine an anxiety phenotype requires some careful consideration. In behavioral pharmacology, construct validity assumes that different assays measuring the same latent trait will yield convergent results (Belzung and Griebel, 2001). Because the mouse OF and EPM converged solely on general locomotion in this cohort, a single OF session may not fully capture trait anxiety. Center time in a novel OF test might more closely reflect locomotor reactivity to the new environment. Therefore, our finding suggests that an anxiolytic-like effect in the OF does not necessarily predict a similar behavioral response in the EPM. Pharmacological data support this. In a systematic review of 814 mouse studies, the benzodiazepine diazepam had a significant effect on EPM open-arm time in 84% of comparisons, but on OF center time in only 59% and on OF distance traveled in 18% (Rosso et al., 2022). Across the seventeen test measures examined, only EPM open-arm time and light– dark box light-compartment time detected anxiolytic effects reliably overall, and no OF measure did. Indeed, critical evaluations of unconditioned models have raised concerns about the empirical evidence supporting the validity of the OF and EPM as interchangeable measures of a unitary anxiety trait (Ennaceur, 2014).

Several limitations should be considered. First, a limitation of this study is that each behavioral test was only conducted once, preventing us from calculating standard test-retest reliability. The activity measures correlated across the two testing environments, indicating that they were captured consistently, but prior work has shown that center time and open-arm exploration are less consistent across repeated testing (Tucker and McCabe, 2017). Second, the mice were group housed and while this is recommended from an animal welfare standpoint the establishment of social hierarchies within cages can introduce individual variability in baseline stress and anxiety-like behavior. Third, the estrous cycle of the female cohort was not tracked, and hormonal phase may interact with both activity and threat appraisal. Finally, our findings come from a single strain and a single cohort. The specific correlations reported here should not be assumed to generalize to other strains, ages or lighting conditions, although the broader observation that the two tests converge on activity rather than on anxiety is consistent with prior literature on cross-test correlation.

In conclusion, the OF and EPM converge on locomotor activity but not on anxiety-like behavior. Activity measures correlated across the two tests, survived correction for multiple comparisons and clustered together, whereas OF center exploration was unrelated to every EPM measure of open-arm exploration and formed a cluster of its own. Sex differences were absent over the 15-min cumulative OF session and, in the EPM, took the form of greater closed-arm activity in females without any difference in open-arm exploration. Furthermore, the rate at which activity habituated within the OF differed between the sexes. Center exploration in a novel OF should therefore not be treated as interchangeable with open-arm avoidance in the EPM, and a single novel OF session may measure locomotor reactivity to novelty rather than the anxiety trait that it is often believed to measure.

## Acknowledgments

During the preparation of this manuscript, the authors used Claude (Anthropic) to assist with editing text, with writing code for the statistical analyses and figures. All analyses of variance were independently run and verified in SPSS by the authors, who reviewed and edited all content and take full responsibility for the accuracy and integrity of the manuscript.

## CRediT authorship contribution statement

**Guido Huisman:** Investigation, Project administration, Formal analysis, Writing – review & editing.

**Lara Caglayan:** Investigation, Project administration, Writing – review & editing.

**Marcelo Febo:** Resources, Supervision, Writing – review & editing

**Chengguo Xing:** Conceptualization, Supervision, Project administration, Writing – review & editing, Funding acquisition.

**Adriaan W. Bruijnzeel:** Conceptualization, Supervision, Formal analysis, Writing – Original Draft, Visualization, Project administration, Funding acquisition.

## Data Integrity and Sponsor Role

The authors confirm that they have had full access to all the data in the study and take responsibility for the integrity of the data and the accuracy of the data analysis. No sponsor was involved in the study design, data collection, or writing of the manuscript.

## Design and Analysis Transparency

The datasets generated and analyzed during the current study are available from the corresponding author on request.

