## Supplement for "Convergent activity but divergent anxiety dimensions in male and female C57BL/6 mice across the open field and elevated plus maze"

**Supplementary tables**

**Supplementary Table 1. Open field behavior, mean ± SEM.**

| **Measure** | **Window** | **Male (n = 29)** | **Female (n = 32)** | **p** |
| --- | --- | --- | --- | --- |
| Center time (% of session) | 15 min | 38.81 ± 3.79 | 42.92 ± 4.33 | 0.481 |
|  | 0–5 min | 35.77 ± 4.28 | 40.67 ± 4.34 | 0.426 |
|  | 5–10 min | 41.48 ± 3.97 | 43.91 ± 4.83 | 0.702 |
|  | 10–15 min | 39.16 ± 3.75 | 44.17 ± 4.35 | 0.391 |
| Center distance (% of total) | 15 min | 47.02 ± 2.30 | 49.50 ± 3.07 | 0.526 |
|  | 0–5 min | 43.09 ± 3.05 | 46.82 ± 3.19 | 0.404 |
|  | 5–10 min | 50.05 ± 2.34 | 52.38 ± 3.38 | 0.581 |
|  | 10–15 min | 49.86 ± 1.85 | 51.34 ± 3.00 | 0.683 |
| Total distance (cm) | 15 min | 2461.1 ± 142.5 | 2409.5 ± 132.6 | 0.792 |
|  | 0–5 min | 946.9 ± 62.5 | 1019.6 ± 60.5 | 0.407 |
|  | 5–10 min | 818.0 ± 51.0 | 732.3 ± 47.4 | 0.223 |
|  | 10–15 min | 696.1 ± 43.9 | 657.5 ± 42.2 | 0.529 |
| Stereotypy counts | 15 min | 2004.0 ± 85.6 | 1858.9 ± 72.2 | 0.197 |
|  | 0–5 min | 664.0 ± 45.4 | 731.7 ± 32.6 | 0.224 |
|  | 5–10 min | 733.8 ± 31.9 | 616.6 ± 33.9 | **0.015** |
|  | 10–15 min | 606.2 ± 27.8 | 510.5 ± 25.3 | **0.013** |

P values from a one-way ANOVA with sex as a between-subjects factor. The whole 15 min session and the three consecutive 5 min blocks are shown. Values are mean ± SEM.

**Supplementary Table 2. Elevated plus-maze behavior, mean ± SEM.**

| **Measure** | **Male (n = 29)** | **Female (n = 32)** | **p** |
| --- | --- | --- | --- |
| Open-arm entries (n) | 8.55 ± 0.81 | 8.22 ± 0.61 | 0.741 |
| Closed-arm entries (n) | 15.93 ± 0.85 | 19.97 ± 0.90 | **0.002** |
| Center entries (n) | 19.34 ± 1.25 | 20.53 ± 1.22 | 0.500 |
| Open-arm entries (% of arm entries) | 34.02 ± 2.41 | 28.82 ± 1.51 | **0.067** |
| Open-arm duration (s) | 61.50 ± 6.55 | 57.11 ± 4.44 | 0.575 |
| Closed-arm duration (s) | 184.79 ± 8.96 | 193.60 ± 5.04 | 0.384 |
| Time on open arms (% of arm time) | 25.67 ± 2.86 | 22.85 ± 1.77 | 0.395 |
| Total distance (cm) | 1005.7 ± 54.5 | 1133.1 ± 45.8 | **0.077** |

P values from a one-way ANOVA with sex as a between-subjects factor. A single 5 min trial. Values are mean ± SEM.

**Supplementary figures**

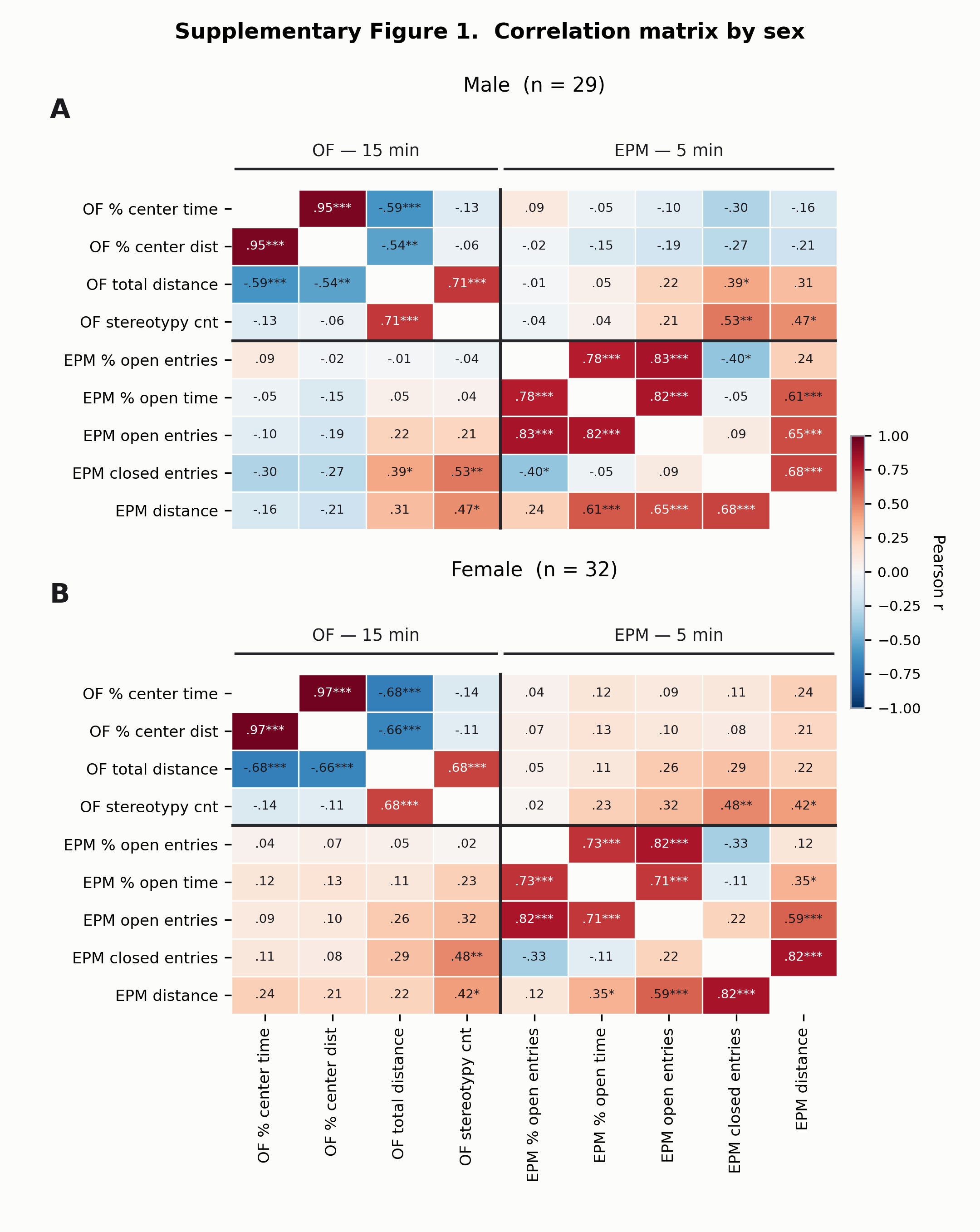

**Supplementary Figure 1. The dissociation between convergent activity and divergent anxiety measures persists across both sexes.** The matrix of Figure 4 computed separately for males (A; n = 29) and females (B; n = 32). The same pattern is present in both sexes: activity pairs correlate, anxiety pairs do not. No cell survives Holm correction at this sample size and none is therefore outlined. Cells show Pearson correlation coefficients; the color scale runs from r = −1 (blue) through r = 0 (white) to r = +1 (red). Asterisks denote uncorrected two-tailed p values: * p < 0.05, ** p < 0.01, *** p < 0.001. Bold outlines mark cells that survive Holm correction.

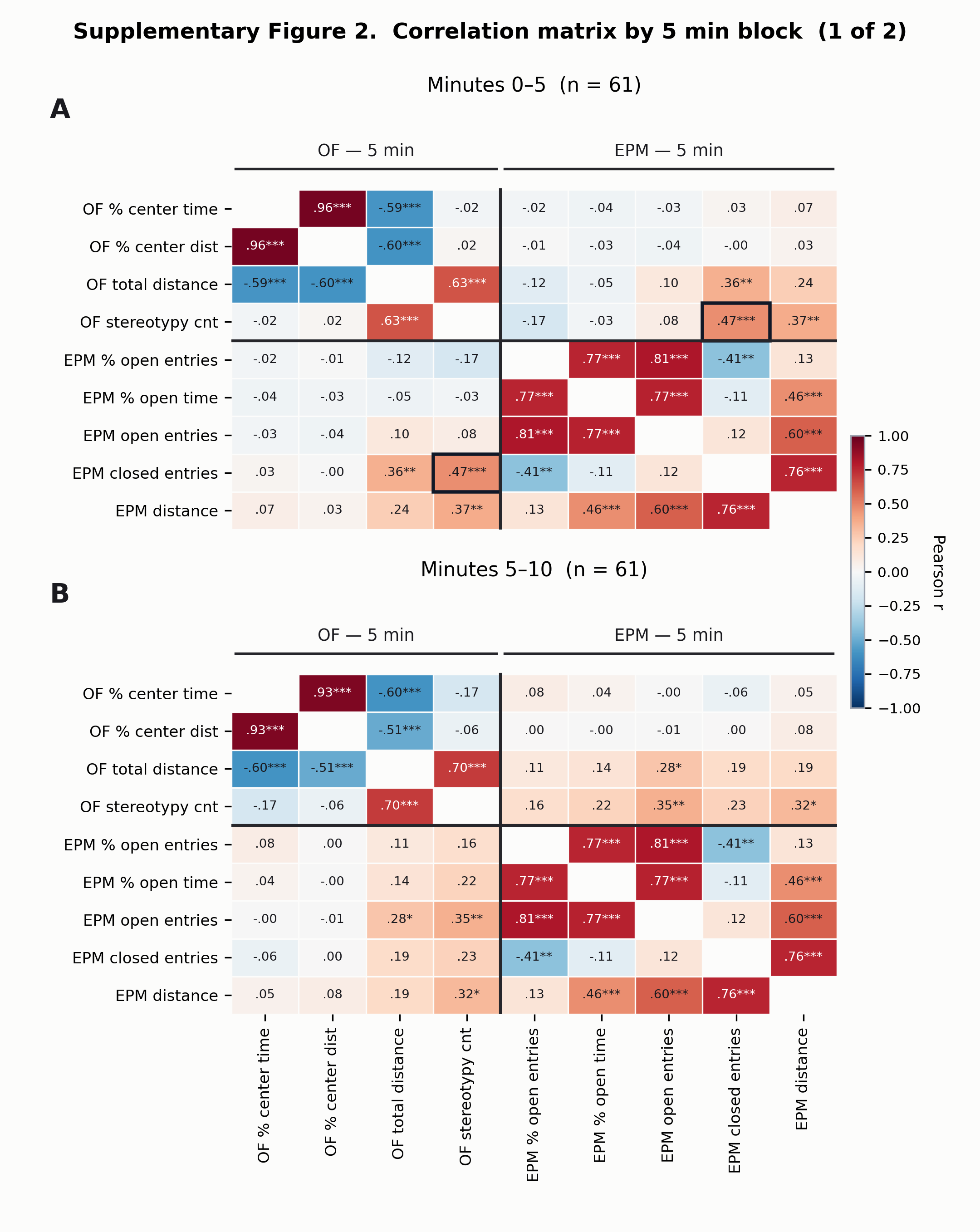

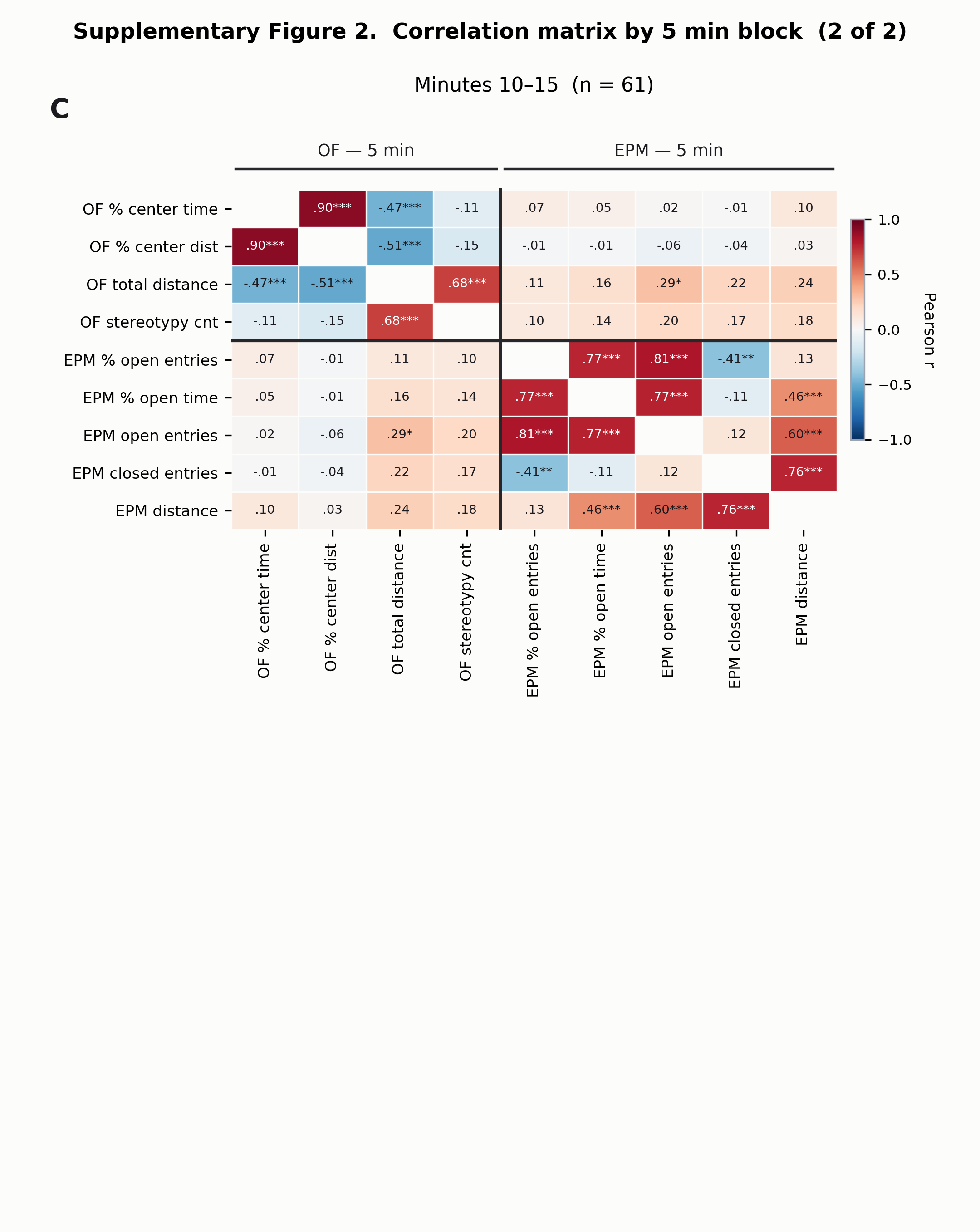

**Supplementary Figure 2. Cross-test correlation matrices demonstrate the temporal weakening of activity coupling across 5-minute blocks.** Correlation matrices computed from minutes 0–5 (A), 5–10 (B) and 10–15 (C) of the open-field session in the same 61 animals, against the same plus-maze trial. The activity coupling is strongest in the first block and weakens across the session. The anxiety pairs remain at zero throughout the three 5-min blocks. Cells show Pearson correlation coefficients; the color scale runs from r = −1 (blue) through r = 0 (white) to r = +1 (red). Asterisks denote uncorrected two-tailed p values: * p < 0.05, ** p < 0.01, *** p < 0.001. Bold outlines mark cells that survive Holm correction.
